# Protein Language Models Uncover Carbohydrate-Active Enzyme Function in Metagenomics

**DOI:** 10.1101/2023.10.23.563620

**Authors:** Kumar Thurimella, Ahmed M. T. Mohamed, Daniel B. Graham, Róisín M. Owens, Sabina Leanti La Rosa, Damian R. Plichta, Sergio Bacallado, Ramnik J. Xavier

**Affiliations:** Broad Institute of MIT and Harvard, Cambridge, MA, USA; Center for Computational and Integrative Biology and Department of Molecular Biology, Massachusetts General Hospital, Harvard Medical School, Boston, MA, USA; Department of Chemical Engineering and Biotechnology, University of Cambridge, Cambridge, UK; School of Medicine, University of Colorado Anschutz Medical Campus, Aurora, CO, USA; Faculty of Chemistry, Biotechnology and Food Science, Norwegian University of Life Sciences, Ås, Norway; Department of Pure Mathematics and Mathematical Statistics, University of Cambridge, Cambridge, UK

## Abstract

In metagenomics, the pool of uncharacterized microbial enzymes presents a challenge for functional annotation. Among these, carbohydrate-active enzymes (CAZymes) stand out due to their pivotal roles in various biological processes related to host health and nutrition. Here, we present CAZyLingua, the first tool that harnesses protein language model embeddings to build a deep learning framework that facilitates the annotation of CAZymes in metagenomic datasets. Our benchmarking results showed on average a higher F1 score (reflecting an average of precision and recall) on the annotated genomes of *Bacteroides thetaiotaomicron*, *Eggerthella lenta* and *Ruminococcus gnavus* compared to the traditional sequence homology-based method in dbCAN2. We applied our tool to a paired mother/infant longitudinal dataset and revealed unannotated CAZymes linked to microbial development during infancy. When applied to metagenomic datasets derived from patients affected by fibrosis-prone diseases such as Crohn’s disease and IgG4-related disease, CAZyLingua uncovered CAZymes associated with disease and healthy states. In each of these metagenomic catalogs, CAZyLingua discovered new annotations that were previously overlooked by traditional sequence homology tools. Overall, the deep learning model CAZyLingua can be applied in combination with existing tools to unravel intricate CAZyme evolutionary profiles and patterns, contributing to a more comprehensive understanding of microbial metabolic dynamics.

## Introduction

Rapid advancements in sequencing technologies have led to an abundance of genomic data, outpacing the capacity to annotate and decipher the functions of these sequences^1^. A significant challenge arises in contextualizing the vast number of unknown functions present in microbes^2,3^ and, as a consequence, a substantial fraction of microbial proteins remains unannotated^4–6^. The Unified Human Gastrointestinal Protein (UHGP) catalog alone holds greater than 170 million protein sequences of which 40% lack any functional annotation^2^. Elucidating the function of these sequences has the potential to provide insights into microbial metabolic behaviors and niches within a particular ecosystem, including the dynamics of microbial-host interactions^7–10^.

In microbial genomics, accurate annotations of the biological functions of enzymes is critical, as these molecules have important roles in catalyzing essential biochemical reactions with high specificity and efficiency^11–14^. Carbohydrate-active enzymes (CAZymes) play fundamental roles in various biological processes, including cell structure, signaling, energy storage, and nutrient processing^15–17^. Metagenomic sequencing and functional ‘omics have shown that CAZymes support the growth of beneficial microbes in infants by catabolizing human milk oligosaccharides (HMOs)^18,19^. CAZymes have also been found to play a role in the microbiomes of patients with inflammatory diseases like Crohn’s disease (CD)^20^ and IgG4-related disease (IgG4-RD), in which there is upregulation of glycan-related pathways^21^.

Historically, functional annotation tools have relied on hidden Markov models (HMMs)^22,23^ that are built by aligning many amino acid sequences or using sequence homology tools like BLAST, which employs a pairwise alignment strategy between query and target sequences^24,25^. The current state-of-the-art tool for annotating CAZymes, dbCAN2, similarly relies on sequence homology or HMMs^26^. While having achieved significant effectiveness in genomic sciences, these methods are not able to assign a biological role to one-third of all bacterial proteins^27^. Advancements in deep learning have significantly aided the functional annotation of proteins and comprehension of their diverse functions^28–35^. Protein language models (pLMs), such as those used for structural prediction and other tasks, demonstrate remarkable capabilities in decoding the intricate amino acid language of proteins, which facilitates their functional annotation through a distinct approach compared to sequence-based alignment methods^30,36–39^. CAZymes are classified into distinct classes of glycoside hydrolases (GHs), polysaccharide lyases (PLs), glycosyltransferases (GTs) and carbohydrate esterases (CEs). Within a class, the enzymes share a conserved fold, mechanism, and catalytic residues^16^. With this fine grained ontology and a set of distinct enzymatic reactions, CAZymes represent an ideal training dataset for pLMs.

Here, we present CAZyLingua, the first annotation tool to harness pLMs for the accurate classification of CAZymes. We applied CAZyLingua to gene catalogs derived from human microbiome metagenomic datasets and identified CAZymes implicated in health and disease states. Our first gene catalog was constructed from paired mother/infant metagenomes^40^ consisting of ∼2,000,000 proteins from which we uncovered ∼27,000 CAZymes previously undetected by dbCAN2 or eggNOG. Early persistence of diverse microbial strains in the gut has been linked with metabolic pathways utilizing CAZymes, including breakdown of HMOs and dietary polysaccharides and metabolism of mucin in the colon^41^. CAZyLingua was then applied to a metagenomic dataset derived from patients with inflammatory and fibrosis-prone diseases, including CD and IgG4-RD. We observed that a greater percentage of genes significantly less abundant in CD were predicted to be CAZymes, while in IgG4-RD, we found an expansion of hundreds of CEs in particular. We demonstrate that CAZyLingua achieves high model accuracy compared to standard sequence homology tools and can be used to augment the functional annotation of CAZymes in metagenomic studies, providing valuable insights into the diversity and functional potential of these microbial enzymes.

## Results

### CAZyLingua Model and Performance

The CAZyLingua pipeline consists of multiple components (Figure 1a). First, the pLM ProtT5^38^ is used to generate embeddings for a given query of amino acid sequences. Second, a quadratic discriminant analysis (QDA) classifier^42^, which takes as an input the ProtT5 embedding, is applied to predict whether the query is a CAZyme or not. Finally, if the query is predicted to be a CAZyme, a multiclass classifier is used to make an annotation in the CAZy database ontology, returning either a family or subfamily. The multiclass classifier was built to return probabilities associated with the given family or subfamily annotation and can return a top *k* number of family labels for a given protein sequence.

**Figure 1.**
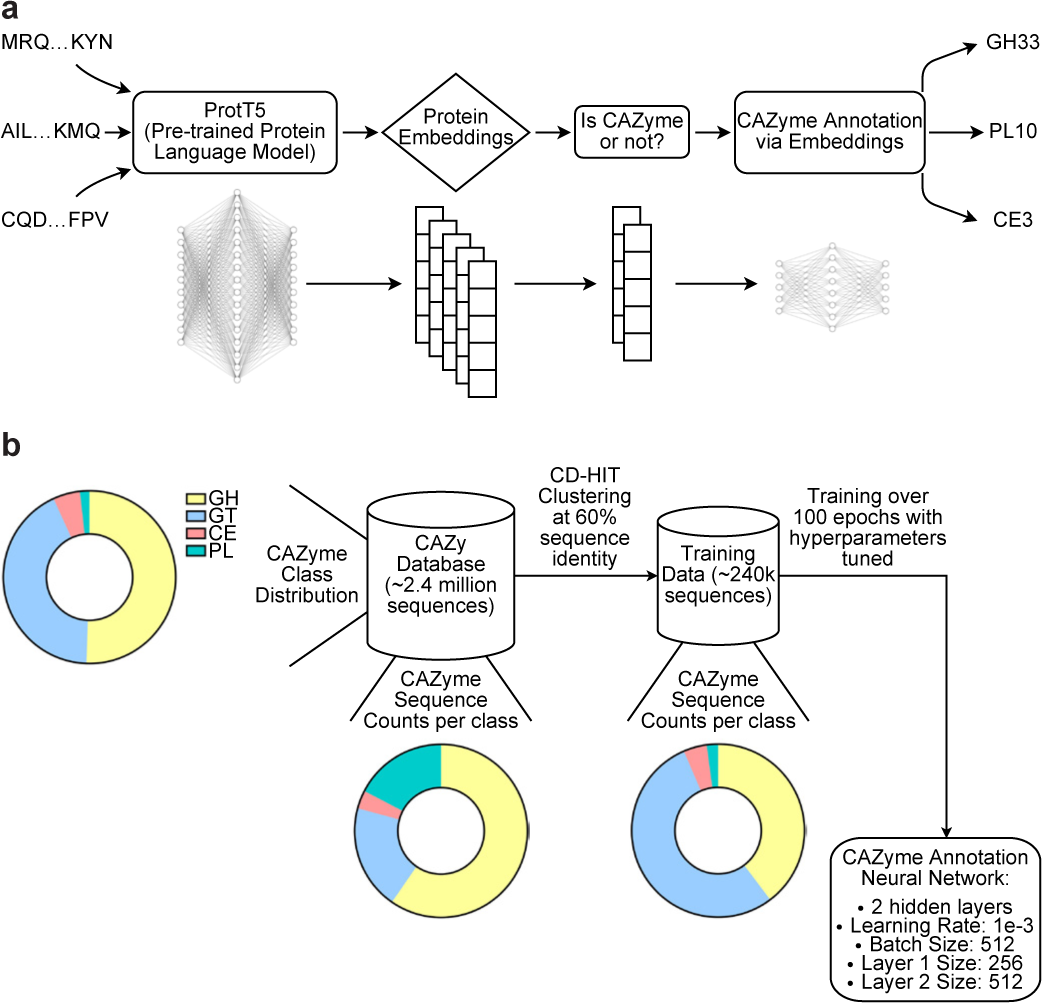
CAZyLingua: a deep learning model used for the classification of proteins as CAZymes. **a)** The workflow of CAZyLingua starts with raw embeddings from ProtT5 followed by the use of those embeddings as input through two classifiers to distinguish 1) whether the embedding was a CAZyme and if so, 2) to which CAZyme family it belongs to. **b)** The training strategy for CAZyLingua began with a 60% sequence identity clustering to remove redundancy from the CAZy database in order to train on distinct CAZymes. The Cross Entropy loss function was applied for training and the loss function that was used included a weighted balancing function to proportionally sample the number of representative sequences per CAZyme class/family/subfamily in the database. This strategy was employed so as not to oversample on highly represented families.

We trained CAZyLingua on a subset of the CAZy database^16,43^ (Figure 1b). CAZymes were selected from every family, spanning GHs, GTs, PLs, and CEs, to create a representative training dataset. To benchmark our method, we followed a procedure similar to dbCAN2, the current state-of-the-art automated CAZyme annotation tool in the community^26^. We specifically chose the DIAMOND+CAZy option in dbCAN2 as this was the closest representation to BLASTp sequence homology. We performed a taxonomic split on the original CAZy database sequences and selected 3 bacterial genomes with pre-annotated CAZymes in each genome: *Bacteroides thetaiotaomicron, Eggerthella lenta,* and *Ruminococcus gnavus*. We selected these bacteria based on the varying proportions of CAZymes per number of total proteins (*B. thetaiotaomicron*: 7.6%, *E. lenta*: 1.1%, and *R. gnavus*: 3.0%) as well as biological relevance: *E. lenta* is very prevalent and found in the gut microbiomes of 80% of humans^44^, *R. gnavus* is linked to patients with CD and produces a proinflammatory carbohydrate^45^, and *B. thetaiotaomicron* is one of the most prevalent members of the gut microbiota and dedicates a large portion of its genome to the processing and utilization of carbohydrates^46^. We obtained these exact protein sequences from the CAZy sequence database to use as the reference set for dbCAN2 DIAMOND+CAZy.

We ran the protein sequences through dbCAN2 and CAZyLingua and evaluated the binary classification task of detecting whether the protein is a CAZyme or not. We combined the results and stratified them into three sets based on whether the protein was predicted by dbCAN2 only, CAZyLingua only, or both. The precision was calculated as the number of true positives in each set divided by the number of predictions made in each set, and recall was calculated as the true positives in each set divided by the total number of CAZymes in each genome (Figure 2a). CAZyLingua alone performed better than dbCAN2 in each measure, but the best benchmarks were in the set of proteins predicted by both tools. We then calculated the F1 score as the harmonic mean of the precision and recall and demonstrated that CAZyLingua outperformed dbCAN2 on each test genome, notably by almost 10% for *E. lenta* (Figure 2b). We examined the predictions by CAZyLingua based on CAZy classes and observed that CAZyLingua was able to label all CE and GT classes in the test genomes (Figure 2c). We evaluated the precision/recall and ROC curves for CAZyLingua and dbCAN2, comparing the output of the decision function from the QDA and the e-value from dbCAN2. Our results showed that CAZyLingua can detect up to 92% of the CAZymes while maintaining a precision of over 80%, while dbCAN2 can detect approximately 82% of the CAZymes at the same precision threshold. CAZyLingua has a higher true positive rate compared to dbCAN2 for this current benchmark (Figure 2d).

**Figure 2.**
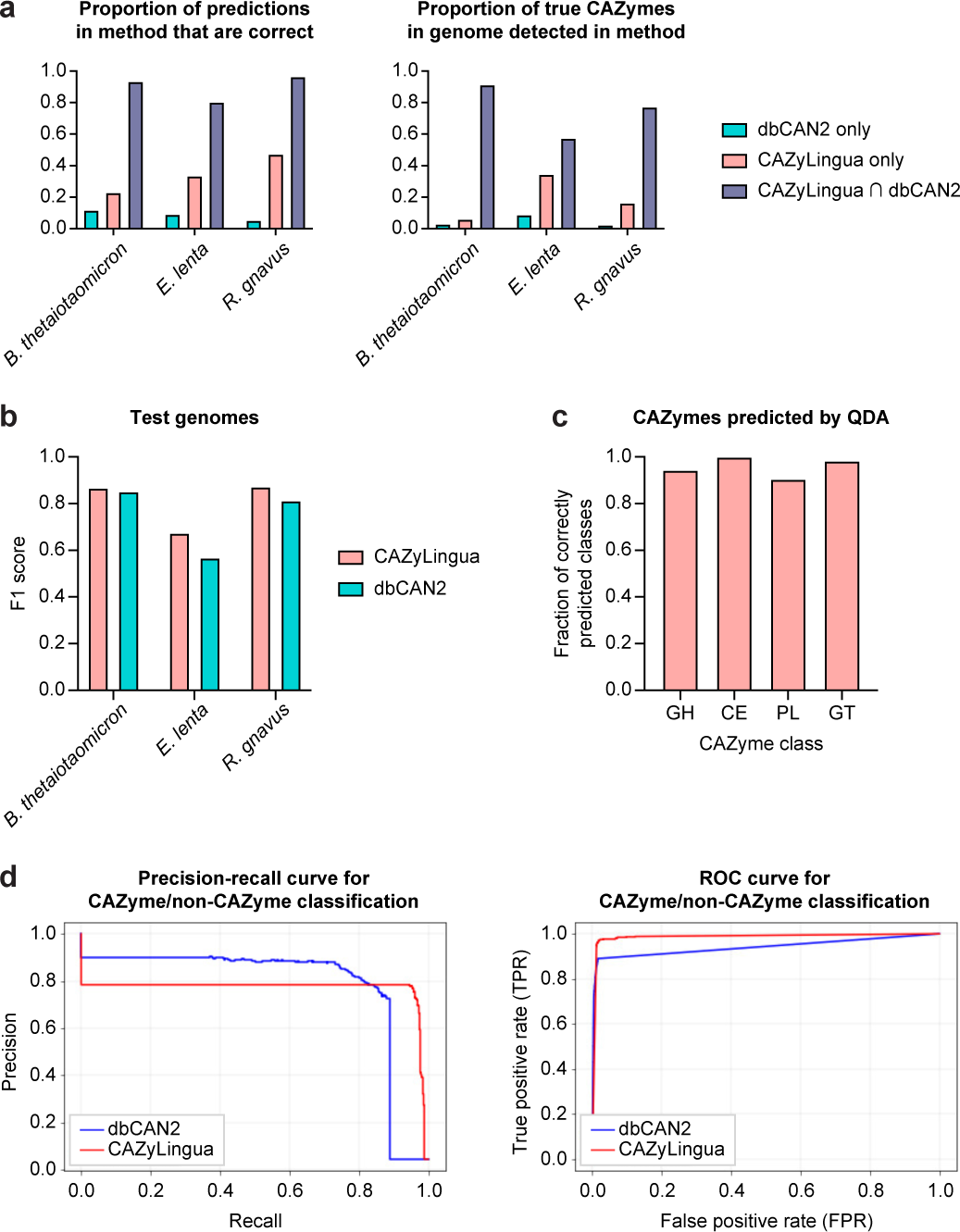
CAZyLingua performance relative to the BLAST-based CAZyme annotation tool dbCAN2. CAZyLingua was compared to the dbCAN2 DIAMOND+CAZy annotation tool option (benchmarked with an e-value < 1×10^-102^). A similar procedure as dbCAN2 was followed by picking 3 bacterial strains with manual annotations and varying CAZyme counts per strain. **a)** For predictions by CAZyLingua only, dbCAN2 only, and shared between the two methods, the proportion of correct predictions made by each method (left) and the proportion of true CAZymes made by each method (right) were calculated. **b)** F1 scores (harmonic means of precision and recall) of all CAZyLingua predictions, all dbCAN2 predictions, and all predictions combined, whether shared between the methods or not. **c)** Ground truth CAZymes were stratified by class, and the percentage of accurate predictions per CAZy class from our Quadratic Discriminant Analysis (QDA) binary classifier was calculated. **d)** Precision/recall (left) and ROC (right) curves comparing CAZyLingua to dbCAN2. The output of the decision function of the boundary that was trained for CAZyLingua and the e-value for dbCAN2 were used for target scores.

For the CAZyme family classification step, we trained over the entire dataset more than 100 epochs, using RayTune^47^ to select different random hyperparameter settings and the best of 20 different training models. The models were all trained with a cross-entropy loss, and RayTune was optimized to store the model on a metric to minimize loss^48^. The best performing model (lowest loss value) was saved, with the corresponding hyperparameter configuration for any CAZyme family inference. The CAZyme classifier is a four-layer, feedforward neural network (with two hidden layers) with an input of 1024 dimensions (fixed size from ProtT5 embeddings) projected to 256 dimensions then to 512 dimensions to a final classification output layer of 574 corresponding to all the unique CAZyme families and subfamilies in our training dataset. We used a hyperbolic tangent (Tanh) as the non-linearity between the different layers. After training, the weights between the first and second layers do not correspond to any interpretable features in the embedding itself (Extended Data Figure 1). When checking a micro-averaged classification accuracy of all the families in the test genomes, CAZyLingua predicted 99.6% of the families accurately, while dbCAN2 predicted 98.2% accurately.

### CAZyLingua Identifies Horizontally-Transferred Genes as CAZymes

We further tested if CAZyLingua would be able to uncover CAZymes in a gene catalog of microbiome samples from mother-infant pairs collected from late pregnancy to one year of age^40^. We predicted CAZymes using CAZyLingua, alongside eggNOG and dbCAN2, on the entirety of the gene catalog, which contained 2,327,970 genes. CAZyLingua predicted 81,498 CAZymes, while dbCAN2 and eggNOG predicted 77,614 and 38,862 CAZymes, respectively. We stratified the dataset by number of genes per sample, then by sample month, and split the observations by mother and infant. We calculated the fold change between each method and CAZyLingua based on the genes per sample per month to determine how many more CAZymes were predicted by CAZyLingua. CAZyLingua predicted at least 2-fold more new genes in maternal and infant metagenomes compared to eggNOG and on average 1.2-fold more new genes than dbCAN2 (Figure 3a). When examining the predictions made by CAZyLingua, we observed 27,133 unique CAZyme predictions that were not made by dbCAN2. We distinguished each unique CAZyme by CAZyme class within each sample over each sample month. We observed that our model predicted many more GTs across all the samples in every month (Figure 3b).

**Figure 3.**
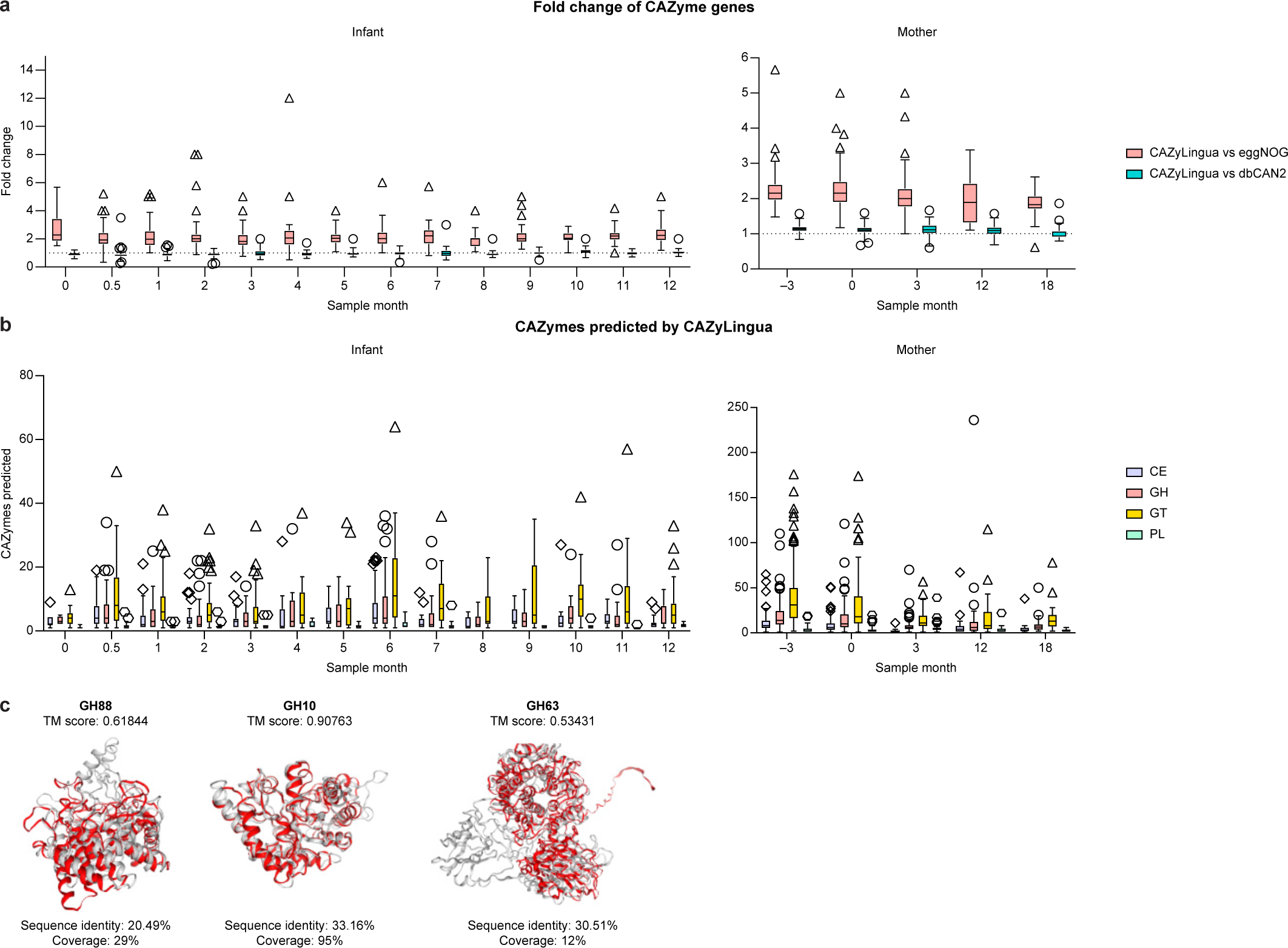
Application of CAZyLingua to metagenomes in paired mothers and infants. **a)** Comparison of CAZyLingua to eggNOG and dbCAN2 on a large metagenomics gene catalog from mothers and their infants. Time of the sample is in months relative to childbirth (month 0). Dotted lines represent no fold change. **b)** CAZyLingua predicted 27,133 genes that dbCAN2 did not, shown by CAZy class for all infant and maternal samples at each sample month. Boxplots in **a** and **b** show medians and interquartile ranges (IQRs), with whiskers showing ± 1.5 IQR. **c)** Predicted structures of proteins from CAZyLingua (red) and the protein embedding nearest neighbor (grey) structurally aligned with TM scores, and BLAST metrics for GH88, GH10, and GH63.

We next focused on a subset of the metagenomic data to specifically look at genes that were found to be horizontally transferred between a mother/infant pair. A previous study performed a sequence homology (BLASTn) analysis on DNA sequences between maternal and infant metagenomes and identified 977 genes with 100% nucleotide identity that were harbored by both maternal and infant species^40^, a portion of which were predicted to function in carbohydrate metabolism. Of the 977 genes, 12 were predicted as CAZymes by our model and either not predicted or predicted as an unknown family within a CAZyme class by dbCAN2.

In order to understand the structural contributions of language models to the general predictions given from ProtT5 and ultimately our pLM classifier, we searched for nearest neighbors between our 12 horizontally-transferred gene embeddings in the CAZy database embeddings using Euclidean distance. After identifying nearest neighbor pairs and extracting the corresponding protein sequences, we computed structural predictions for those proteins using ColabFold^49^. We used FoldSeek^50^ to perform a structural alignment between the structures of the predicted protein from CAZyLingua and the nearest protein embedding neighbor in the CAZy database.

CAZyLingua predicted four GHs, including three belonging to the families 88, 10, and 63, that had a high structural homology to their nearest neighbor in the CAZy database (all with a TM score > 0.50, which indicates a same fold between two proteins^51^). In contrast, when evaluating sequence homology (BLASTp) between the amino acid sequences of the three proteins and the nearest neighbor in the CAZy database, we found that between both sets of sequences the sequence identity was lower than 35%, and for GH88 and GH63 the coverage was less than 30% (Figure 3c). Given these metrics, this suggests that CAZyLingua is able to predict CAZymes incorporating structural homology, despite the lack of any amino acid sequence homology.

The fourth GH predicted was given the annotation of GH43_18 when evaluating the ProtT5 nearest neighbor, while CAZyLingua classified it as a GH33 (Figure 4a). We sought to explain if the classification of a GH33 was based on specific features of the unknown CAZyme. We first evaluated the neighborhood of genes around the unknown CAZyme to establish if it exists in a functional polysaccharide utilization locus (PUL). We found several canonical PUL features, including several regulatory elements related to carbohydrate metabolism: a hybrid two-component system (HTCS), TonB-dependent receptor (SusC homolog), and contiguous substrate-binding lipoprotein (SusD homolog) (Figure 4b). In addition to this unknown enzyme mapping to a PUL, we established the presence of a lipoprotein signal peptide in the enzyme through SignalP^52^. We then explored the link between several functional sites in the GH33 and the corresponding embedding generated by ProtT5. To do so, we created a sliding window of 10 amino acids and created more distant substitutions of the original sequence within that window based on the BLOSUM62 distance. Substituting areas near the signal peptide corresponded to the greatest losses in the CAZyLingua predictive value of a GH33. The first 20 amino acids that correspond to a signal peptide were used in a homology search, and in all BLAST metrics, the signal peptide showed stronger homology to GH33: a combined percent identity and coverage of 64.2% for GH33 and 55.0% for GH43_18, providing stronger evidence for its classification as a GH33 (Figure 4b).

**Figure 4.**
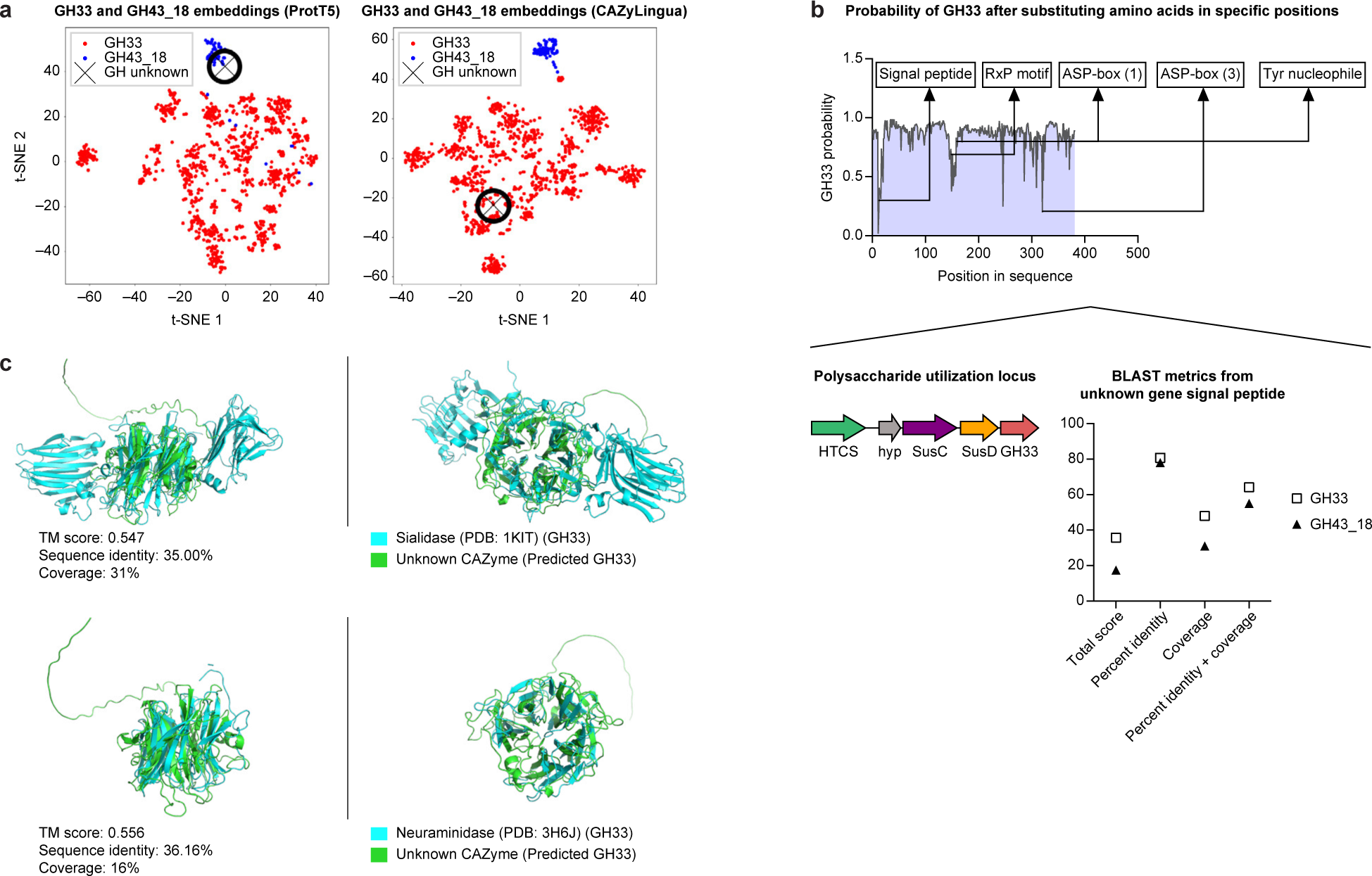
CAZyLingua distinguishes GH33 CAZyme from nearest neighbors of raw ProtT5 embeddings. **a)** tSNE of (left) ProtT5 embeddings from the GH33 and GH43_18 families and the CAZyme predicted by CAZyLingua (GH unknown) and (right) a segment of the last layer of CAZyLingua. **b)** GH33 protein residues were mutated in a sliding window of ten residues over the entire sequence, and ProtT5 embeddings were generated for each sliding window mutation. Known features are overlaid along sections of the sequence. The probability of the CAZyLingua-predicted classification being a GH33 was calculated for each sliding window mutation (top). The predicted GH mapped to a PUL containing several regulatory elements consistent with a CAZyme (bottom left). BLAST metrics on the predicted GH signal peptide compared with GH33 and GH43_18 sequences (bottom right). **c)** Overlays of the predicted GH protein structure generated using ColabFold with a sialidase (top) and a neuraminidase (bottom).

To determine if there was any structural homology between our unknown CAZyme and the GH33 family, we used ColabFold^49^ to fold our protein and ran a structural search with 3D crystal structures found in the PDB25 database using DALI^53^. Our unknown protein had several matches, with two in the top five matches being GH33-like enzymes, namely a neuraminidase and a sialidase. After structurally aligning^51^ our unknown structure with the neuraminidase and the sialidase crystal structures, we observed that the predicted GH33 shared significant structural homology (TM score > 0.5) with both. The sequence homology (BLASTp) between the amino acid sequences pairwise with the unknown protein revealed sequence identities <36% and coverages <31% (Figure 4c).

### Analysis of Enriched CAZymes in Inflammatory Disease Metagenomic Gene Catalogs

We next focused our attention on applying CAZyLingua to two metagenomic datasets derived from patients with inflammatory and fibrosis-prone diseases: one from 68 CD patients and 34 control subjects ^54^ and another from 58 IgG4-RD patients and 165 healthy controls^21^. Both of these disease states have unique microbial signatures potentially underlying pathologic mechanisms.

To investigate disease-associated genes that may be unannotated CAZymes, we first used a linear model against the CD gene catalog^55,56^ (Methods) and identified 3,499 genes that were significantly more abundant (two-sided *t*-test, p < 1×10^-2^, log fold change > 2) and 30,125 genes that were significantly less abundant (two-sided *t*-test, p < 1×10^-2^, log fold change < −2) in CD. Among these, CAZyLingua predicted 30 more abundant genes and 569 less abundant genes to be CAZymes (Figure 5a, Supplementary Table 1). Given the ∼10-fold difference between more abundant genes in controls versus CD, we observe many more glycan-related pathways associated with health compared to CD.

**Figure 5.**
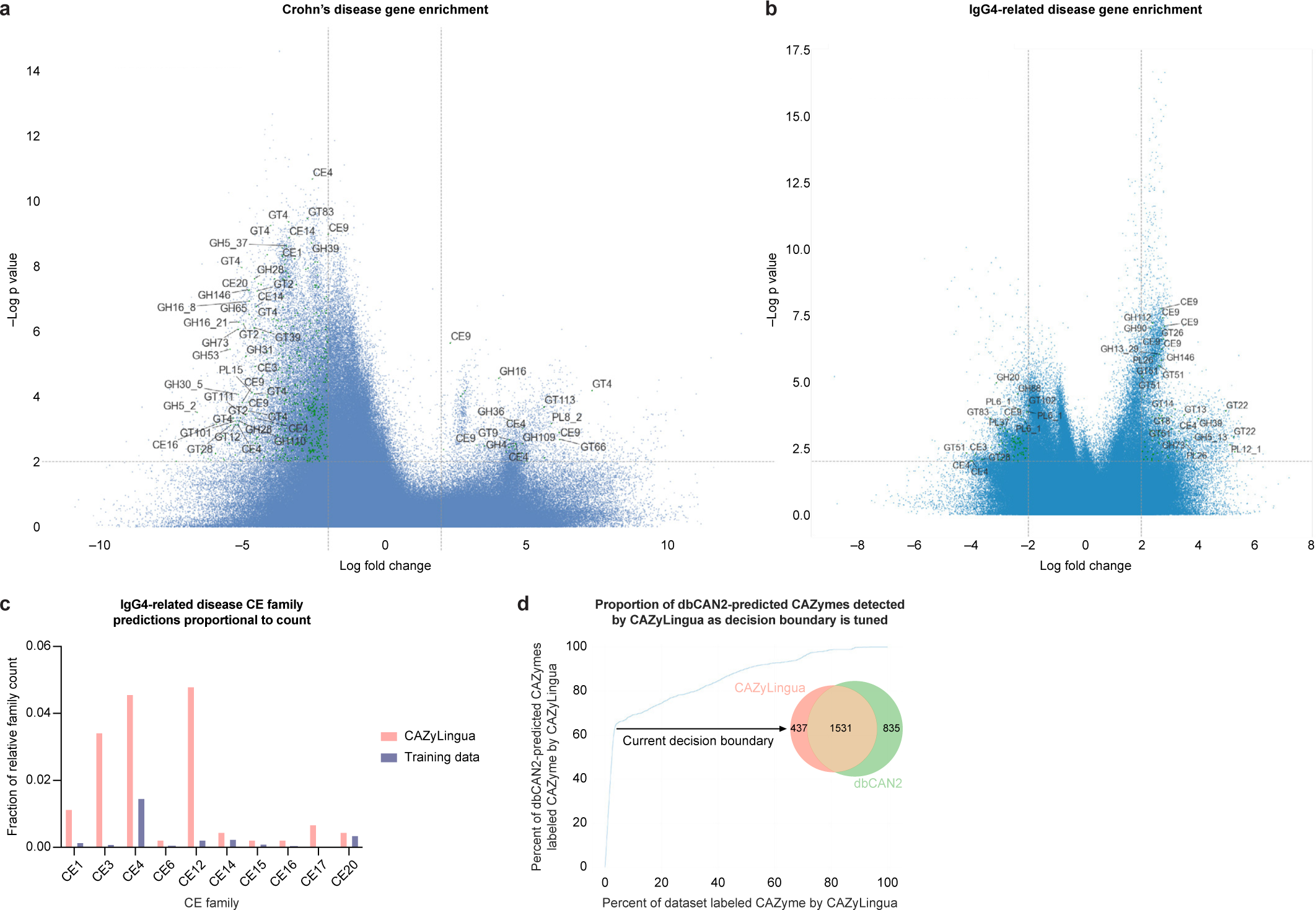
Application of CAZyLingua to CAZymes in metagenomes of patients with inflammatory and fibrosis-prone diseases. Genes enriched and depleted in the gene catalogs of patients with **a)** CD and **b)** IgG4-RD selected on the fringe of the volcano plot (see Methods for labeling criteria). **c)** Predicted CEs in the enriched IgG4-RD gene set, stratified to analyze only the genes CAZyLingua predicted. **d)** The proportion of dbCAN2-predicted CAZymes also predicted by CAZyLingua as the decision function between CAZyme/non-CAZyme of the QDA classifier in CAZyLingua was varied. The Venn diagram shows the numbers of CAZymes predicted by CAZyLingua, dbCAN2, and both on our current model benchmarks of the QDA.

Following the same analysis procedure, we built a linear model for a differential gene abundance analysis for IgG4-RD metagenomes. We stratified genes based on the same criteria. Compared with the CD dataset, we noticed a higher proportion of genes were significantly more abundant in IgG4-RD compared to a healthy state. We observed 9,225 genes that were significantly more abundant compared to 7,284 genes that were significantly less abundant in IgG4-RD. CAZyLingua predicted 65 more abundant and 87 significantly less abundant CAZymes in IgG4-RD (Figure 5b, Supplementary Table 2).

We then broadened our focus to all the CAZymes in the IgG4-RD dataset, irrespective of their significance to disease from the linear model. CAZyLingua predicted 437 CAZymes that dbCAN2 did not. Specifically in IgG4-RD, there was a higher number of CEs that only CAZyLingua predicted. CE sequences comprise only 4% of all the sequences in the CAZy database; the low representation of certain sequence examples can pose a challenge for sequence homology tools, which may explain the lower number of hits identified by dbCAN2. In our set of genes predicted by CAZyLingua only, we observed that ∼34% were CEs. Families of CEs that were particularly represented included CE1, CE3, CE4, and CE12 (Figure 5c). All of these families share SGNH (Ser-Gly-Asn-His) hydrolase activity, which is a conserved structural feature of the enzymes in these families, suggesting that these enzymes may have low sequence homology but higher structural homology within each class^57–59^.

The increase in annotations by CAZyLingua for these specific CE families may be due to the unique structural features of the families that otherwise would be hard to annotate by traditional sequence homology methods. Given the distinct set of CAZyme families that CAZyLingua was able to predict, we sought to determine the extent of overlap between CAZyLingua predictions and the set of CAZymes that dbCAN2 annotated. To learn about the binary classification of CAZyme/non-CAZyme given by the QDA predictions and the results from dbCAN2, we varied the QDA decision boundary. We calculated the percentage of CAZymes that CAZyLingua labeled as CAZyme that dbCAN2 also predicted against the percentage of the entire IgG4-RD gene set that CAZyLingua labeled as CAZyme. Our QDA model was benchmarked where ∼5% of the dataset was labeled CAZyme by CAZyLingua and that represents ∼60% of all the genes that dbCAN2 also predicted as CAZyme. At ∼30% of the dataset being labeled as CAZyme by CAZyLingua, we captured ∼80% of all the dbCAN2-predicted CAZymes. As we relaxed our decision boundary and increased the number of genes in the dataset CAZyLingua labeled as CAZyme, we observed a relatively linear relationship between the genes labeled as CAZyme by both dbCAN2 and CAZyLingua (Figure 5d). This linear relationship describes a relative discordance between the annotations from the two different tools. The divergence of annotations generated by CAZyLingua compared to dbCAN2 can add to existing CAZyme annotations in the analysis of large metagenomics studies.

## Discussion

In this study, we introduced CAZyLingua, a novel approach that leverages pLMs to enhance the identification and functional annotation of CAZymes in metagenomic datasets. Our method mitigates the ongoing challenge of assigning functions to the vast array of unannotated microbial enzymes within these datasets, shedding light on their potential roles in various biological processes. The use of pLMs has emerged as a powerful tool for unraveling protein functions in microbial genomics^28–30^, and our results further emphasize their efficacy in this context. When compared with traditional sequence homology, CAZyLingua improved the F1 score of classifying a protein as a CAZyme by 6.1% for each of the benchmarked test genomes with gold standard annotations.

CAZyLingua’s efficacy is evident in its successful identification of previously undiscovered CAZymes within a longitudinal microbiome dataset of mother-infant pairs. We detected over 27,000 unique putative CAZymes that were missed by dbCAN2. Furthermore, our identification of horizontally-transferred CAZymes between mothers and infants highlights the ability of CAZyLingua to uncover potentially crucial enzymatic functions that traditional sequence homology methods might overlook. When investigating GHs that were missed by dbCAN2, we noticed that these GH structures shared low sequence homology (sequence identity < 40%) to the most homologous protein in the embedding latent space. Our analysis of structural similarities between CAZyLingua-predicted enzymes and GH structures highlights the potential of CAZyLingua to predict enzyme functions based on structural conservation (TM score > 0.5), thereby offering insights into their catalytic roles. We note that these findings are based on structural predictions from ColabFold, not crystal structures or experimentally validated enzymes. One advantage to our choice of ColabFold as a structural prediction tool is that the process of generating a prediction is heavily dependent on a multiple sequence alignment (MSA) between an unknown sequence and a large reference of sequences. The goal of using ColabFold over popular pLM-based structural prediction tools (e.g., ESM-fold, OmegaFold) was for there to be less of a bias between predictions based on embeddings in a process similar to CAZyLingua and how ProtT5 may be trained versus a standard MSA.

We focus on an example of a horizontally-transferred GH33 that was not predicted by dbCAN2, eggNOG, or a nearest neighbors search using ProtT5 in the CAZy database. Upon using ColabFold to fold this GH33, we performed a sensitive structural search using DALI^53^ against experimentally-characterized crystal structures (PDB25) and found the top hits to include other GH33 enzymes (a sialidase/neuraminidase), with significant structural homology (TM score > 0.5, Z score > 2). A recent study examining the early colonization of microbes in a murine model^60^ highlights an example of vertical transmission of a GH33 sialidase (NanH) between dams and pups. The NanH gene is triggered by sialylated host glycans and aids in the early colonization of *Bacteroides fragilis*. The putative GH33 discovered by CAZyLingua that was transmitted between a maternal *Alistipes finegoldii* strain and an infant *Alistipes putredinis* strain might exhibit similar properties as NanH and could be part of a mechanism to aid in the establishment of *Alistipes putredinis* in the infant gut. Again, sequence homology between our putative GH33 and NanH was low (33.93% identity, 26% coverage) despite a similar predicted function, indicating that existing sequence homology methods might have overlooked the putative GH33 as a functional homolog. This highlights the strengths of pLMs as alternative tools to augment functional protein homology discovery.

We then extended the utility of CAZyLingua to metagenomic datasets from patients with CD and IgG4-RD. Both diseases share pathological features of fibrotic lesions despite having distinct clinical presentations. Patients with CD have been shown to have lower microbial diversity and carbohydrate utilization pathways in their gut microbiota^61–63^. Unique microbial signatures have been strongly associated with IgG4-RD, and those signatures included genes linked to carbohydrate metabolism^21^. Our initial analysis focused on genes that were upregulated in IgG4-RD, where we found a distinct set of CAZymes using CAZyLingua. Investigating the taxonomy of those genes, we found several from *Streptococcus* species that are typically found in the oral cavity. In the previous study^21^, many *Clostridium* and typically oral *Streptococcus* species were overabundant in the disease phenotype while *Alistipes* and *Bacteroides* species were depleted. Six of the top 20 (30%) putative CAZymes predicted by CAZyLingua mapped to *Streptococcus mutans,* and we observed that many genes from this microbe were upregulated in disease. We observed enrichment of CEs within this species and postulated that there may be several CAZymes that help *Streptococcus mutans* adapt to an ecological niche in the gastrointestinal tract of patients with IgG4-RD.

CEs themselves were sparsely populated in our training dataset for CAZyLingua and similarly in the CAZy database of sequences. Due to the imbalance of this class of enzymes, we postulate that sequence homology may fail to annotate these enzymes. During our training procedure, we use a weighted cross entropy loss, where the weights are proportional to the number of training examples for a given CAZyme family or subfamily. By allowing a more stringent penalty on incorrectly annotating a rare family, we are able to predict more rare families like CEs through CAZyLingua.

The implications of our findings extend beyond the specific datasets analyzed in this study. CAZyLingua’s demonstrated ability to accurately predict CAZymes has broader implications for deciphering the functional potential of microbial communities. A similar procedure of fine-tuning pLM embeddings can be broadly applied to other enzyme classes and protein domains to aid in functional annotation. As an ever-growing number of metagenomic datasets become available, the incorporation of deep learning tools like CAZyLingua into existing methods offers a promising avenue for comprehensive and accurate functional annotation.

## Methods

### CAZyme training dataset curation

The CAZy database found at http://www.cazy.org/IMG/cazy_data/cazy_data.zip is cataloged by the dbCAN tool maintainers and a fasta file is available at https://bcb.unl.edu/dbCAN2/download/. We downloaded the CAZy database as of August 06, 2022 containing 2,428,817 sequences as it was the latest version that was available for when we began training the model. We chose to focus on the four main classes CAZymes: 173 families and 177 subfamilies in glycoside hydrolases (GHs), 115 families in glycosyltransferases (GTs), 20 families in carbohydrate esterases (CEs), and 42 families and 60 subfamilies in polysaccharide lyases (PLs). We removed everything that did not belong to one of these families and any sequences that were larger than 5000 amino acids in length to prevent GPU out of memory errors when generating embeddings. The entire number of remaining sequences was 2,413,796: 1,221,013 in GH, 1,027,247 in GT, 122,413 in CE, and 43,123 in PL.

Using the CD-HIT software tool^64^, we clustered our CAZy database at 60% sequence identity. CD-HIT returns a representative sequence for a given cluster. The clusters were created such that, in the resulting database (nr.CAZy.60.fasta), no two sequences had a sequence similarity greater than 60%. The resulting database preserved all of the original families and subfamilies while reducing the redundancy in the database. The database in nr.CAZy.60.fasta contained 232,736 sequences, of which 92,385 sequences were in GH, 125,240 in GT, 10,177 in CE, and 4,934 in PL.

Following the curation of the CAZy sequences, we used ProtT5^38^ to generate embeddings for each of these sequences using a V100 GPU. We stored the embeddings in h5 files, following the hierarchical data format (HDF). This embedding database served as the training dataset for both of the classifiers in CAZyLingua.

### Quadratic discriminant analysis training and testing

To build the CAZyme/non-CAZyme binary classification step in the CAZyLingua pipeline we modeled the embeddings from the CAZy training dataset as our positive case (CAZyme) and used a combination of data from protein families database Pfam and the Kyoto Encyclopedia of Genes and Genomes (KEGG) to construct our negative examples (non-CAZyme). We started with the 1,296,280 Pfam seeds as a dataset from which to construct negative examples. Pfam seeds serve as the basis for hidden Markov model (HMM) profiles and are highly curated to span a diversity of domains^65^. This dataset has been previously described as building the HMMs that contribute to greater than 75% of all the functional annotations of Uniprot sequences in Pfam^28^. We additionally supplemented the negative examples with 3,435 enzymes from KEGG that were non-CAZymes using the KEGG Enzyme database^66^.

In order to create a set of negatives on which to train, we used the ultra-sensitive parameter of DIAMOND^67^ in the BLASTp setting between the Pfam seeds against the CAZy database and then the KEGG enzymes against the CAZy database. We removed any Pfam seeds or KEGG enzymes that were listed as hits from the DIAMOND output. The remaining 56,244 Pfam seeds and 3,429 KEGG non-CAZyme enzymes were combined to create a non-CAZyme dataset. We sampled 5,000 CAZymes from nr.CAZy.60.fasta spanning all families and subfamilies in each class as our positive example.

We built our model using scikit-learn^42^, importing the function QuadraticDiscriminantAnalysis with the store_covariance parameter selected as true. We used the library skops to pickle and save the state of our trained model. For a given set of embeddings, the QDA classifier will label them as CAZyme or non-CAZyme and store the results of the CAZy embeddings in an h5 file.

To model our QDA, we model the distribution of each embedding whether it is a CAZyme or not a CAZyme. These form our two classes *c*: CAZyme and non-CAZyme.

We model the prior probability of a class *c*, *P*(*y* = *c*) by the empirical proportion of training samples in that class. The conditional probability of a protein’s embedding *x* ∈ ℝ^1024^ given its class *c*, *P*(*x* | *y* = *c*) is modeled by a multivariate Gaussian distribution with probability density function:

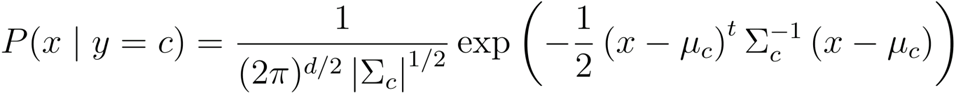

The parameters *μ_c_* and Σ*_c_* for each class are the maximum likelihood estimators given the training samples in that class. If the training samples are (*x_i_*, *y_i_*), *i* = 1, …,*N*, the maximum likelihood estimators are given by

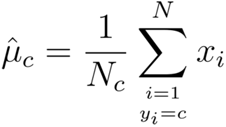

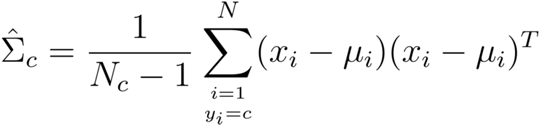

where *N_c_* is the number of samples class *c*.

Predictions for a protein with embedding *x*\* are made by assigning the class *c*\* which maximizes the posterior probability, given by Bayes’ rule:

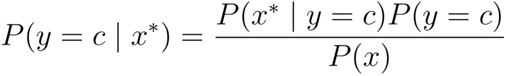

where only the numerator depends on *c*. A decision surface is created for the QDA based on the two classes, CAZyme and non-CAZyme^42^.

In constructing ROC curves, the decision function that we used is the logarithm of the posterior probability.

### Feed forward neural network architecture

The final stage in the CAZyLingua model is the multiclass classification for a given CAZyme family based on the embeddings selected as CAZyme from the QDA. The feedforward neural network architecture has three overall layers with two hidden layers. The fixed size input of 1024 dimensions from ProtT5 embeddings are projected to 256 dimensions then to 512 dimensions to a final classification output layer of 574, which reflects the number of CAZyme families and subfamilies. We implemented this model using Pytorch Lightning^68^ to create a classifier that included all of the training, validation, and testing steps.

The model used a Cross Entropy Loss from PyTorch^48^ with the weights parameter set to balance the number of sequences from the different families and subfamilies. In order to prevent over training on highly represented families, the loss function penalty for a given family was calculated as the inverse of the number of sequences per family. This ensures that if the model is incorrectly labeling a family with very few training examples there will be a stronger penalty in comparison to incorrectly labeling a family with a higher proportion of the training examples.

### Hyperparameter optimization and neural network training

The multiclass classification neural network in the CAZyLingua pipeline was trained using RayTune^47^, a hyperparameter tuning library. The hyperparameters that were tested were the size of layer 1, the size of layer 2, the batch size, and the learning rate. In order to find the optimal hyperparameters to select the most accurately trained model, 20 models were tested in parallel with random sampled hyperparameters selected by RayTune (Supplementary Table 3). Each model was trained over 100 epochs using the Async Successive Halving (ASHA)^69^ scheduler that terminates a model (early stopping) optimized to minimize the training loss. Metrics for the validation accuracy were collected after each epoch, and the testing accuracy was collected after the model was fully trained. Each training model was visualized using TensorBoard^70^ (Extended Data Figure 2).

**Supplementary Table 3.** Hyperparameter Tuning. Training epochs over time to pick the model with the best classification accuracy. Using RayTune, we performed a random grid search of different hyperparameter values and tested 20 models in parallel. We picked the model with the best accuracy and used that as the model for all further inference.

| Hyperparameter | Sampling Method | Sampled Values |
| --- | --- | --- |
| Layer 1 Size | Random Choice | (256, 512, 768) |
| Layer 2 Size | Random Choice | (512, 1024, 1536) |
| Batch Size | Random Choice | (127, 256, 512) |
| Learning Rate | Log Uniform Sample | [1e-4 – 1e-2] |

### Benchmarking of CAZyme/non-CAZyme QDA classifier

To benchmark the QDA classifier, we used different metrics to quantify the performance of CAZyLingua to dbCAN2. For the F1 score, we followed a standard formula:

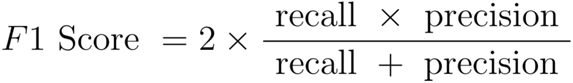

where we define recall and precision as follows:

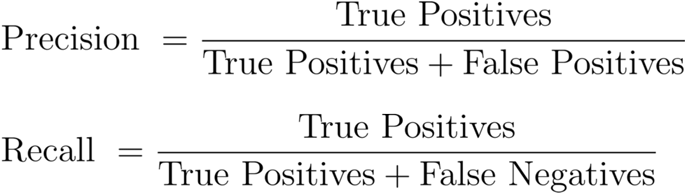

The precision-recall and ROC curves were plotted using sklearn^42^ using the precision_recall_curve and roc_curve using the e-values from dbCAN2 and the scores from the decision function of the QDA from CAZyLingua as the target scores.

We designed two metrics to benchmark the differences between CAZyLingua’s predictions, dbCAN2’s predictions, and the predictions shared by both methods.

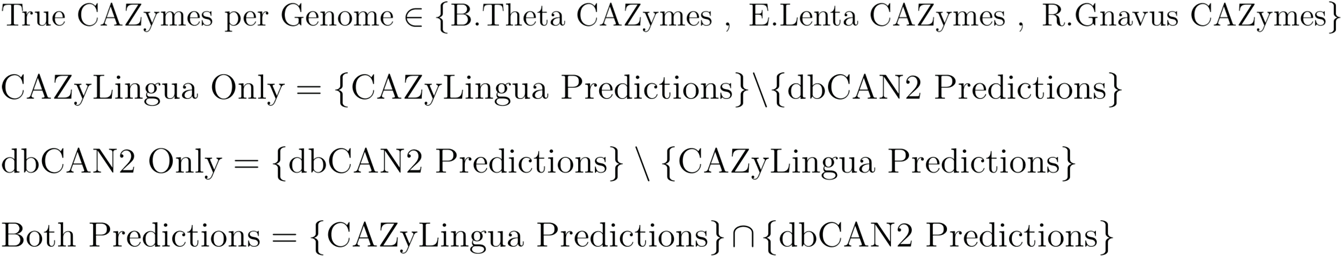

With each of these different sets, we calculated the metric to find the proportion of true CAZymes to all predictions in each genome predicted by each method:

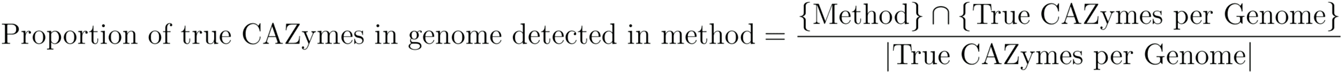

Each method was also benchmarked to find the proportion of annotated CAZymes that were correctly labeled as being CAZymes in each method:

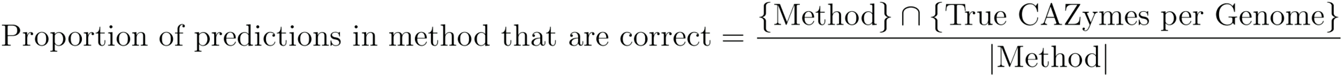

where

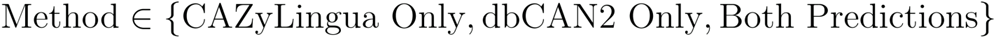

### Gene catalog construction

The metagenomes for each disease type (IgG4-related disease^21^ and Crohn’s disease^54^) and for the mother/infant cohort^40^ were assembled into their respective gene catalogs following the same procedure. A quality control check was performed using Trim Galore!^71^ to remove sequencing adapters and kneadData to remove human reads and trim low quality reads (--trimmomatic-options “HEADCROP:15 SLIDINGWINDOW:1:20 MINLEN:50”) to keep reads that were minimum 50 bp long. All the quality controlled reads were assembled using MEGAHIT^72^. Each contig had all of the open readings frames predicted using Prodigal^73^, and we keep both gene and protein sequences. A non-redundant gene catalog was built with a sequence identity threshold of 95% using CD-HIT^64^. To construct a count matrix, each read was mapped using a Burrows-Wheeler Aligner with at least 95% sequence identity for the length of the read. For determining the taxonomy of each contig, MMseqs2^74^ was used with NCBI RefSeq as the taxonomic annotation database.

The IgG4-RD non-redundant (90% sequence identity) gene catalog consisted of 2,237,319 genes from 58 IgG4-RD samples and 165 healthy control samples^21^. The CD non-redundant (90% sequence identity) gene catalog consisted of 5,929,528 genes from 68 CD samples and 34 non-IBD control samples^54^. The mother/infant non-redundant (95% sequence identity) gene catalog consisted of 2,327,970 genes, with 74 infants, 137 mothers, and 70 mother-infant pairs. Infants were sampled each month between birth (0 months) and 12 months (and additionally at 0.5 months), and mothers were sampled at gestational week 27 (approximately 3 months prior to the birth of the child) and at 3, 6, 9, and 12 months after the birth^40^. Each of these gene catalogs were constructed in each respective prior study and directly utilized in the analysis presented in this paper.

### Analysis of mother/infant gene catalog

The entire mother/infant gene catalog was run through dbCAN2 (diamond blastp -d ${CAZy_reference} -q ${query_file} -o ${output_str}.matches.tsv -e 1e-102 -k 1 -p 2 -f 6) and eggNOG on default parameters. Additionally, embeddings were generated for the entire mother/infant dataset using ProtT5, with CAZyLingua running inference on the entire gene catalog.

We took the 977 horizontally-transferred gene subset and collected all of the dbCAN2 and CAZyLingua results. We took the 12 genes that only CAZyLingua predicted and performed a structural prediction on each of the protein sequences. We performed a Euclidean distance search between those 12 embeddings and the nr.CAZy.60.fasta database to find the closest embedding and subsequently the CAZyme family. We then used ColabFold^49^ to fold each of the 12 proteins and their nearest neighbor to generate PDBs for each horizontally-transferred gene and neighbor pair. A structural alignment was computed on each of these pairs using Foldseek^50^, which returns the overlapped structures and a TM score for each pair. To compute sequence homology metrics, we selected the “Align two or more sequences’’ option in the BLASTp suite on the NCBI website (https://blast.ncbi.nlm.nih.gov/Blast.cgi?PROGRAM=blastp&PAGE_TYPE=BlastSearch&LINK_LOC=blasthome).

The putative GH33 and each of the GH33 and GH43_13 in nr.CAZy.60.fasta were ordinated through tSNE (sklearn TSNE package)^42^ and plotted using matplotlib^75^. A structural prediction of the putative GH33 was produced from ColabFold^49^ and the amino acid residue substitution analysis was done using custom scripts. To search against known, experimentally-characterized structures, the DALI option to pairwise search against PDB25^53^ was used. To structurally align a pairwise hit from putative GH33 to a structure from PDB25, we used US-align^51^ to generate aligned structures and a TM score.

### Disease metagenomic differential abundance analysis

In each disease gene catalog, linear modeling was used to regress different disease covariates onto each gene in the catalog to find differentially abundant genes (features). An abundance filter was applied to the entire count matrix to remove any genes with <10% prevalence across samples. A zero-inflation was applied to any zeros in the count matrix, where the zero value would be replaced by the minimum non-zero value in the given feature and divided by 2. The fold change was calculated by dividing the mean of the disease group by the control group, and taking the log_2_ of the value. Each value is log_2_ transformed and a z-score is calculated for every value in a given feature using the scipy^76^ library. A linear model, from the statsmodels^55^ library, is then applied to each feature. For IgG4-RD, the metadata covariates modeled were: age, on treatment, rituximab, prednisone, other treatments, sex, and cohort. In CD the variables modeled were: age, on antibiotics, mesalamine, and steroids. A significance threshold was established for all of the analyses: we followed a multiple testing adjustment, and p-values were corrected using Benjamini-Hochberg correction, with a false discovery rate (FDR)-corrected p value (q-value) of 0.25. The volcano plots were labeled based on four conditional arguments for the CD and IgG4-RD metagenomic catalogs. For CD, the criteria for the displayed labels were:

1. logFC > 2 and p-value < 1×10^-5^
2. logFC < −2 and p-value < 1×10^-8^
3. logFC > 3 and p-value < 1×10^-2.5^
4. logFC < −4.5 and p-value < 1×10^-3^

For IgG4-RD, the criteria for the displayed labels were:

1. logFC > 2 and p-value < 1×10^-5^
2. logFC < −2 and p-value < 1×10^-3.5^
3. logFC > 3 and p-value < 1×10^-2.5^
4. logFC < −3.5 and p-value < 1×10^-2^

## Acknowledgements

K.T. was supported by the Gates Cambridge Trust and the Rotary Foundation. This work was funded by the National Institutes of Health (DK043351, DK127171, and HL157717 to R.J.X.) and Center for Microbiome Informatics and Therapeutics.

## Extended Data Figure Legends

**Extended Data Figure 1.**
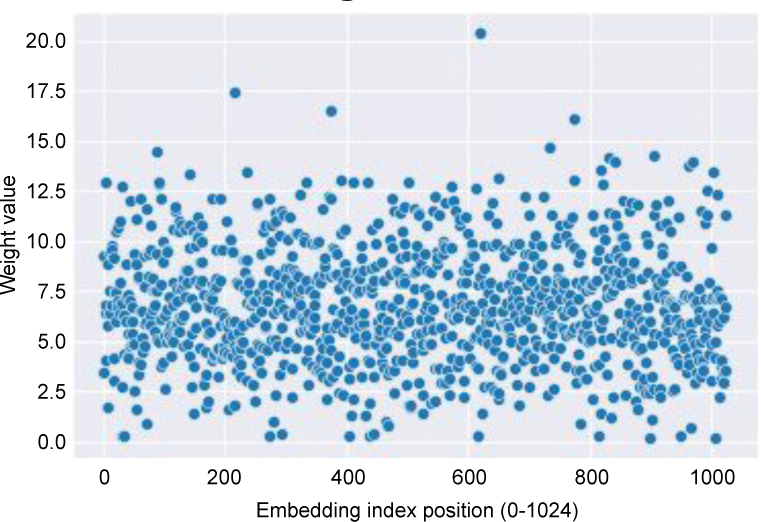
Embedding weights from first layer to next, no interpretable chemical features. We extracted the weights (***W***) from the CAZyLingua multiclass classifier between the input layer and first hidden layer, which is a matrix of dimension 1024×256. After applying a transpose to get ***W***^***T***^we multiplied the two matrices, ***W*** · ***W***^***T***^ which produced a symmetric matrix, ***S*** of dimensions 1024×1024. After taking the ***diag***(***S***) we obtained a vector of size 1024, which is the size of the original embedding from ProtT5. We plotted the values in the vector to visualize if there were any features or positions in specific regions of the embedding that are specific to CAZymes.

**Extended Data Figure 2.**
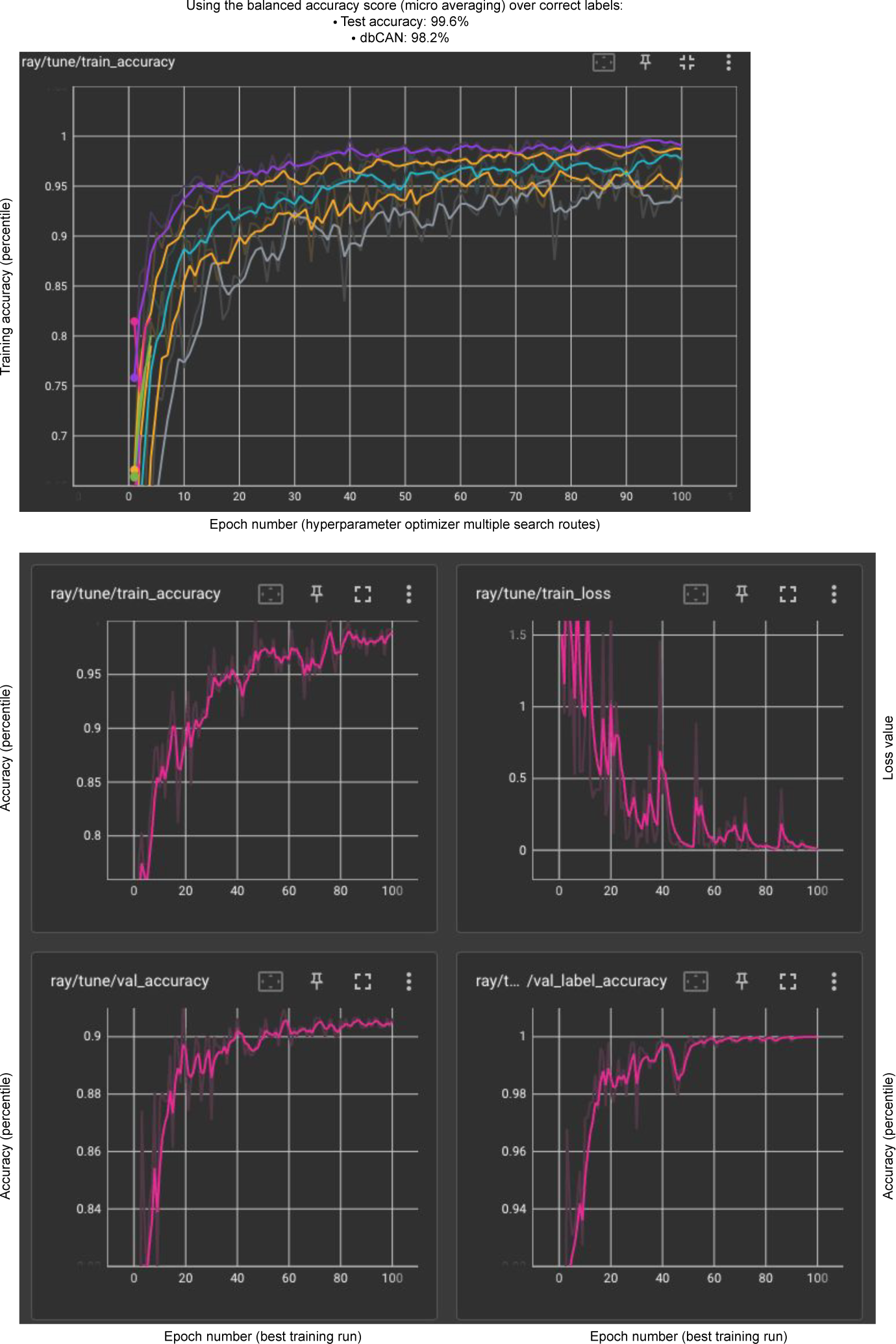
Training runs for finding the best model. RayTune ran 20 models in parallel over each epoch and pruned any models that began to stagnate or have a decline in training accuracy. The models were evaluated on the metric of minimizing training loss, and the model with the minimal loss was stored as a checkpoint. There were 100 epochs over which training occurred, and the metrics were stored and written to a TensorBoard that produced these visualizations.

